# Evaluating Plastic Deformation and Damage as Potential Mechanisms for Tendon Inelasticity using a Reactive Modeling Framework

**DOI:** 10.1101/349530

**Authors:** Babak N. Safa, Andrea H. Lee, Michael H. Santare, Dawn M. Elliott

## Abstract

Inelastic behaviors, such as softening, a progressive decrease in modulus before failure, occur in tendon andare important aspect in degeneration and tendinopathy. These inelastic behaviors are generally attributed to two potential mechanisms: plastic deformation and damage. However, it is not clear which is primarily responsible.In this study, we evaluated these potential mechanisms of tendon inelasticity by using a recently developed reactive in elasticity model (RIE), which is a structurally-inspired continuum mechanics frame work that models tissue in elasticity based on the molecular bond kinetics. Using RIE, we formulated two material models, one specific to plastic deformation and the other to damage. The models were independently fit to published experimental tensile tests of rat tail tendons. We quantified the inelastic effects and compared the performance of the two models in fitting the mechanical response during loading, relaxation, unloading, and reloading phases. Additionally, we validated the models by using the resulting fit parameters to predict an independent set of experimental stress-strain curves from ramp-to-failure tests. Overall, the models were both successful in fitting the experiments and predicting the validation data. However, the results did not strongly favor one mechanism over the other. As a result, to distinguish between plastic deformation and damage, different experimental protocols will be needed. Nevertheless, these findings suggest the potential of RIE as a comprehensive framework for studying tendon inelastic behaviors.

## 1 Introduction

Inelastic mechanical behaviors, where energy is dissipated during mechanical loading, occur during tendon’s overload and in loading-induced pathological conditions such as degeneration and tendinopathy [1,2]. Inelastic behaviors in tissue include viscoelasticity, plastic deformation, and damage. In particular, plastic deformation and damage are associated with stress softening, commonly referred to as Mullin’s effect in polymer mechanics [3,4]. In tissue, softening is observed as a progressive decrease in tensile modulus prior to failure [5–8] and can be described based on the concepts of plastic deformation and damage. In plastic deformation, there is a shift in the unloaded reference state, and in damage the material loses its ability to absorb energy without a change to the unloaded reference state [3,9,10]. These inelastic behaviors can contribute to the development of clinical disorders, such as tendinopathy and rupture by changing the macro-scale mechanical behavior [11] and can also affect tissue degeneration and regeneration, by changing the cell loading [12].

Tendon’s mechanical and structural studies support both of the potential mechanisms of plastic deformation [13–15] and damage [16,17] for softening. However, plastic deformation and damage are hard to differentiate experimentally. Plastic deformation and damage cannot be differentiated during the commonly used ramp-to-failure tests due to the similar softening effect of these two mechanisms during loading, and a repeated loading protocol, such as loading and unloading, is essential to quantify plastic deformation and damage. However, even when unloading is added, the low stiffness in the toe-region and viscoelastic behaviors can still limit differentiation between plastic deformation and damage [14]. Structural studies also do not resolve the question: non-recoverable micro-scale sliding [18] supports plastic deformation, collagen denaturation due to mechanical loading [19,20] supports damage, and the inelastic tensile response of individual collagen fibrils [21,22] may support both. Despite numerous mechanical and structural studies, tendon’s inelastic behaviors are not well described.

We recently developed the reactive inelasticity (RIE) theoretical framework to address the inelastic behaviors of soft tissue by using the kinetics of molecular bonds [23]. This structurally-inspired continuum mechanics framework is an extension of the reactive viscoelasticity model by Ateshian [24] and prior work in polymer mechanics [3,25–27], which addresses all three of the aforementioned inelastic behaviors of tissue (i.e., viscoelasticity, plastic deformation, and damage) by using the same constitutive settings at different kinetic rates of bond breakage and reformation. This characteristic makes RIE distinct from other models for tissue’s inelastic behaviors, which address viscoelasticity without softening [28–30], or include softening by focusing on either plastic deformation [14,31], damage [10,17,32–34], or their combination [35–38]. In the RIE framework there are three bond types; formative bonds are used to model a transient viscoelastic behavior, permanent bonds are used for modeling the hyperelastic behavior, and sliding bonds are used for plastic deformation. Damage is added to each bond type by reducing the number fraction of the available bonds [23]. By combining different bond types, a wide variety of mechanical behaviors can be modeled. Hence, the RIE framework provides a comprehensive tool for investigating the inelastic behaviors of tissue by using a unified structurally-inspired theory to model inelasticity.

Our long-term goal is to study tissue inelasticity, bridging the gap between experiments and modeling. The objectives of this study are to apply the RIE modeling framework to tendon’s tensile behavior, to evaluate the model performance in fitting the experimental data for plastic deformation and damage mechanisms, and to validate the models by predicting an independent experimental dataset. Our previous study on rat tail tendons demonstrated that uniaxial mechanical loading can change both macro-scale mechanical properties and micro-scale structure [18]. However, this experimental study was not designed to distinguish between plastic deformation and damage, and it did not provide numerical measures for inelastic effects. We will show that we can describe the inelastic behaviors in tendon by using RIE-based models specific to plastic deformation (combination of formative bonds and sliding bonds) or damage (combination of formative bonds and permanent bonds). Importantly, for validation we then apply the resulting model parameters to predict results from a separate, independent experiment. This study is significant in that it pursues a systematic approach to elucidate the mechanisms of tissue degeneration and failure, and it provides a path forward for understanding the connection between structural hallmarks of tendon pathology and mechanical behavior.

## 2 Methods

We formulated two material models using the reactive inelasticity (RIE) framework that address either plastic deformation or damage. The models were independently fit to the experimental data from Lee and co-workers [18]. We first evaluated the performance of these two models in fitting the experimental data. To validate the model prediction, we applied the resulting model parameters to a separate, independent experiment that measured the stress response of tendon fascicles in constant-rate tensile tests (Szczesny and co-workers [39]).

### 2.1 Reactive inelastic modeling

In the following, we briefly describe the theoretical formulation of RIE and its specific application to this study; for further details the reader is referred to [23]. The RIE framework employs the kinetics of molecular bonds to simulate the inelastic behaviors. There are two levels for categorizations of bonds: “bond types” and “generations”. A bond type, specifies its mechanical behavior (e.g., hyperelastic or plastic deformation). Bonds break when subjected to external loading, and reform to a new configuration. The reformed bonds initiate a new generation. At any point during loading, each bond type will have several generations with different reference configurations and number fractions that evolve according to a constitutive model for kinetics of molecular bonds. By adding the response of generations, the overall response of a bond type is calculated, and by combining different bond types, an RIE material model is formulated [23].

We used a simplified one-dimensional version of RIE to model the rat tail tendon’s axial tensile tests. We assumed that the stress in the material is the sum of all the bonds types (*T* = Σ_γ_ *T*_γ_), and that the stress of a bond type is a weighted sum of the stresses from generations that depends on the overall stretch λ and the reference stretch of each generation 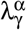, such that 
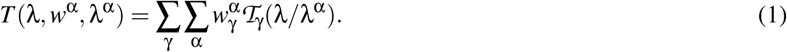

In this equation, *w*^*α*^ is the number fraction of the generation (α) from a bond type (γ), which is a positive scalar less than one, and 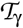 is the intrinsic hyperelasticity stress function that provides a unique stress for each deformation for each generation of a bond type. As a result, the state variables that control stress are λ, 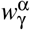, and 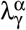 [23].

We modeled the kinetics of molecular bonds to formulate the evolution of 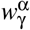, and 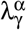 for the different bond types [23]. Three types of bonds are defined based on their kinetic rate: formative (γ = *f*), permanent (γ = *p*), and sliding (γ = *s*), which account for viscoelastic, hyperelastic, and plastic deformation behaviors, respectively [23]. The formative bonds have a finite rate of breakage and reformation and the reference stretch is determined by the deformation at which the generation was initiated. That is, 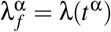, and *t*^α^ is the time at which the generation was generated. This results in a viscoelastic behavior with a stress-free equilibrium condition (Fig. 1A,D) [24,27]. Permanent bonds do not break (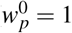 and 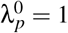), which results in a hyperelastic behavior with no deformation history dependence. The permanent bonds are a special case of formative bonds at the limit of slow kinetics. Sliding bonds are the other limit case; this time the kinetics rate is fast and the bonds almost instantaneously break and reform. In this case, the plastic deformation behavior can be formulated using constitutive relations for 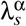, where, for the 1D uniaxial tensile loading, 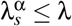 is the sufficient condition [23].

**Fig. 1.**
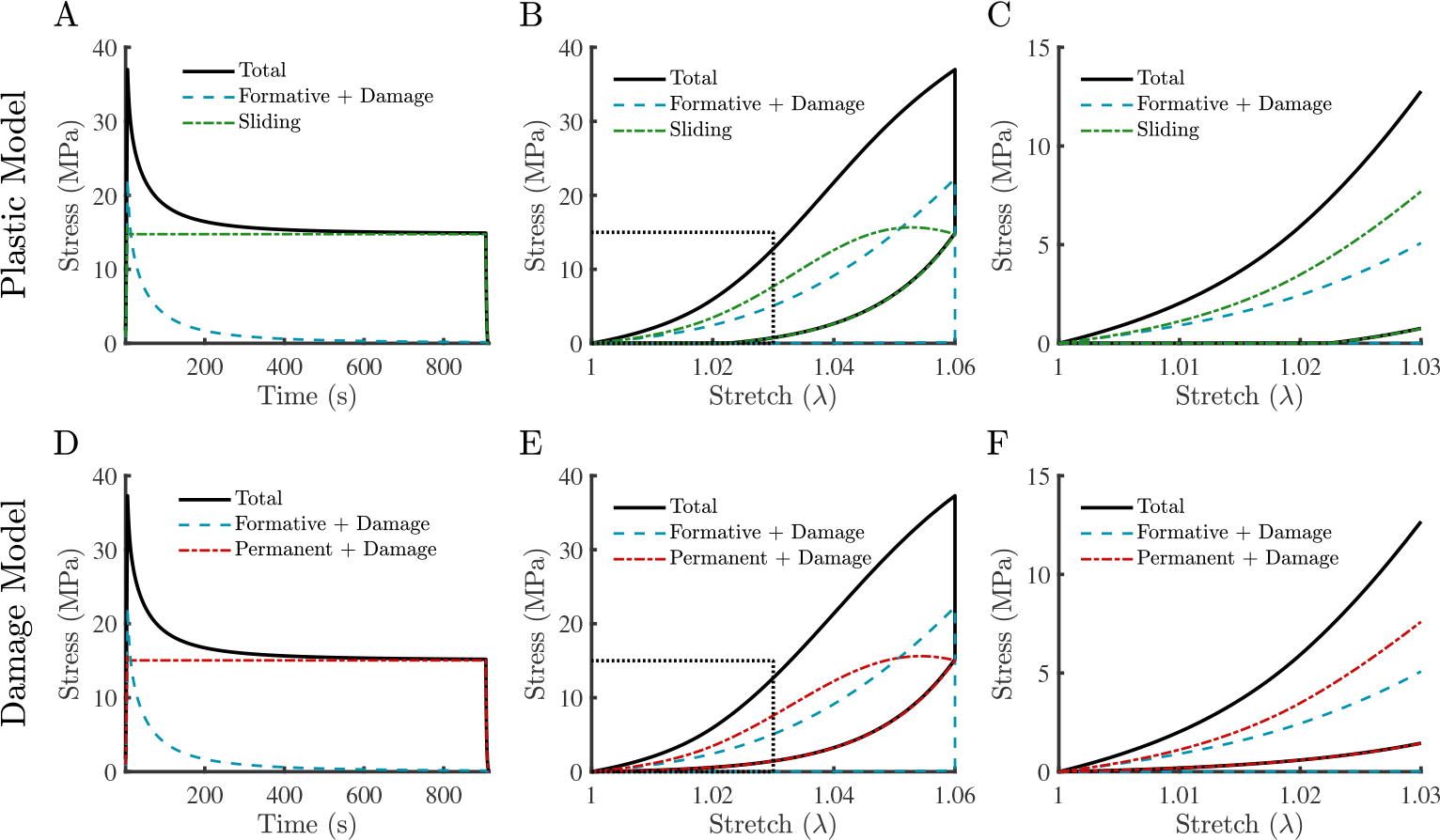
Schematic difference between the plastic deformation and damage models. In the plastic model (A-C) a combination of formative bonds with damage and sliding bonds is used. For the damage model (D-F) a combination of formative bonds with damage, and permanent bonds with damage is used. Both of the models have similar behaviors during loading, and relaxation ((A,D) and (B,E)); however, when looking closer at the boxed region in (B) and (E) during unloading (C and F) the plastic model shows a shift in reference unloaded configuration, where there is none for the damage model.

In this formulation, damage can be applied to all of the bond types by reducing the number fraction of the bonds able to carry load [23]. As a result,

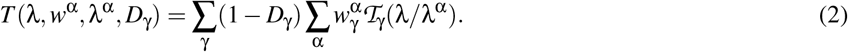

In the above relation, *D*_γ_ is the damage parameter for each bond type that is a scalar in the range [0,1] [34]. Both damage and the change of reference configuration in sliding bonds have a similar behavior during loading (Fig. 1B,E); however, during unloading the difference between these two processes becomes evident. Unlike plastic deformation, damage causes the softening effect without a shift in the reference unloaded configuration (Fig. 1C,F) [23]. Note that for permanent bonds Eq. (2) reduces to

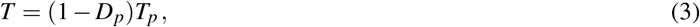

which is the formulation used in classic continuum damage mechanics [34,40,41].

### 2.2 Plastic deformation and damage constitutive relations

In this study, two separate models were created using the RIE framework: (1) plastic deformation model and (2) damage model. For the plastic deformation model we used a combination of formative bonds (γ = *fP*) and sliding bonds (γ = *sP*), where

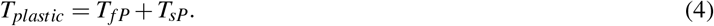

For the damage model, a combination of formative bonds (γ = *fD*) and permanent bonds (γ = *pD*) was used, where

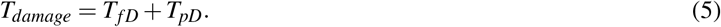

The details of the constitutive relations and parameters are explained in the following and are summarized in Table 1.

**Table 1.** Summary of the bond types, constitutive relations, and the model parameters used for each of the inelastic models.

|  | Bond Type | Kinetics | Intrinsic Hyperelasticity | Sliding | Damage | # of parameters |
| --- | --- | --- | --- | --- | --- | --- |
| **Plastic** | Formative | Nth-order | Fiber-exponential | - | Weibull | 10 |
| | | $[K(s^{-1}), N]_{fP}$ | $[C_1(\text{MPa}), C_2]_{fP}$ | - | $[k, l, r_0]_{fP}$ | |
|  | Sliding | Step | Fiber-exponential | Modified Weibull | - |  |
| | | - | $[C_1(\text{MPa}), C_2]_{sP}$ | $[b, c, r_0]_{sP}$ | - | |
| **Damage** | Formative | Nth-order | Fiber-exponential | - | Weibull | 10 |
| | | $[K(s^{-1}), N]_{fD}$ | $[C_1(\text{MPa}), C_2]_{fD}$ | - | $[k, l, r_0]_{fD}$ | |
|  | Permanent | - | Fiber-exponential | - | Weibull |  |
| | | - | $[C_1(\text{MPa}), C_2]_{pD}$ | - | $[k, l, r_0]_{pD}$ | |

#### 2.2.1 Kinetics

For formative bonds, an nth-order kinetics rate equation was used:

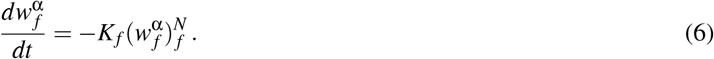

In this relation, the model parameters are (*K*_*f*_, *N*_*f*_), where *K*_*f*_ is a positive number that scales the rate of the reaction and *N*_*f*_ is the order of the breakage reaction (*N*_*f*_ ≥ 1). For the special case of *N*_*f*_ = 1, this relation reduces to a first-order kinetics equation, where its time constant is τ_*f*_ = 1/*K*_*f*_ [23]. For the sliding bonds, a step kinetics relation is used, where for breaking generations 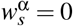, and for reforming generation 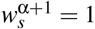 [23]. For the permanent bonds, since there is no bond breakage, τ_*p*_ = 1/*K*_*p*_ → ∞

#### 2.2.2 Intrinsic hyperelasticity

To account for nonlinear stiffening of tissue (toe-region), a typical fiber-exponential constitutive relation was used as the intrinsic hyperelastic function of the bonds [42,43]:

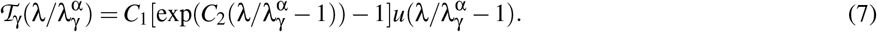

In this equation, (*C*_1_,*C*_2_)_γ_ are positive-valued model parameters. In the plastic model, *C*_1_ and *C*_2_ are the same for formative and sliding bonds, and likewise for the damage model *C*_1_ and *C*_2_ are the same for formative and permanent bonds. This is a simplifying assumption that indicates the bonds from each model have the same load bearing capability, but their difference is due to the difference in their kinetics and inelastic parameters. In addition, *u*(.) is the Heaviside step function, which is included so that the constitutive relation does not provide for stiffness in compression.

#### 2.2.3 Sliding

The axial stretch was used as the sliding variable 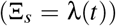, which governs the sliding process in an cumulative way [23]. That is, sliding stretch increases upon loading, and its value does not change during unloading. Hence, the difference between the reference stretch of two consecutive generations of the sliding bonds (α - 1 and α) is calculated as [23]

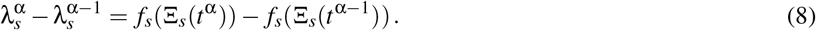

A modified Weibull’s relation was used for *f*_*s*_ such that

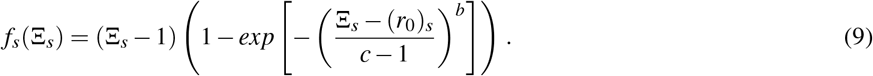

The model parameters are *b* (*b* ≥ 1) the shape parameter, *c* the scale parameter, and (*r*_0_)_*s*_ is the initial threshold of sliding (*c* ≥ 1, (*r*_0_)_*s*_ ≥ 1). For a simple ramp loading with (*r*_0_)_*s*_ = 1, one can show that 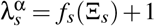. For convenience, the sliding stretch of the last generation 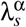 is referred to as the sliding stretch λ_*s*_.

#### 2.2.4 Damage

Similar to the sliding formulation, the axial stretch was used as the damage variable 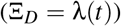 and the Weibull’s cumulative distribution function (CDF) was used for accumulation of damage [33,34]

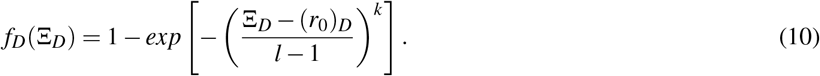

In this equation (*k*, *l*) are the Weibull’s shape and scale parameters, respectively, and (*r*_*D*_)_0_ is stretch at the initial onset of damage (*k* ≥ 1,*l* ≥ 1, (*r*_*D*_)_0_ ≥ 1) [23]. The accumulation of damage is calculated by considering the history of deformation and 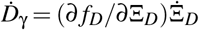 using a non-recoverable scheme that only allows for increase in damage during loading [23]. Similar to 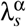 it can be shown that if (*r*_*D*_)_0_ = 1, for a simple ramp loading 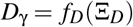. Damage is applied to permanent bonds for the damage model. Additionally, the formative bonds in both models are allowed to accumulate damage.

### 2.3 Experimental micro-tensile test data

Experimental data from Lee and co-workers [18] was used to fit the two models. Briefly, tendon fascicles were dissected from the tail of Sprague-Dawley rats and tested in a PBS bath. The mechanical tensile test consisted of ~ 5 mN preload, 5 cycles of preconditioning to 4% strain, a ramp load at 1%/sec to either 4%, 6%, or 8% grip-to-grip strains (n=7/group)(loading phase), held for 15 minutes (relaxation phase). Following relaxation, the specimen was unloaded at 1%/s (unloading phase), and reloaded to failure (reloading phase) (Fig. 2). The initial cross-sectional area and grip-to-grip length were used to calculate stress and strain. We used the maximum optical strain (reported as the tissue strain in [18]) to scale the grip-to-grip strain to account for the gripping effects and to avoid the errors in estimation of the applied strain to the tissue. The stress data was smoothed using the moving average method and re-sampled to one-thousand data points for each loading, relaxation, unloading, and reloading phases, and were then used for curve-fitting to calculate the model parameters.

**Fig. 2.**
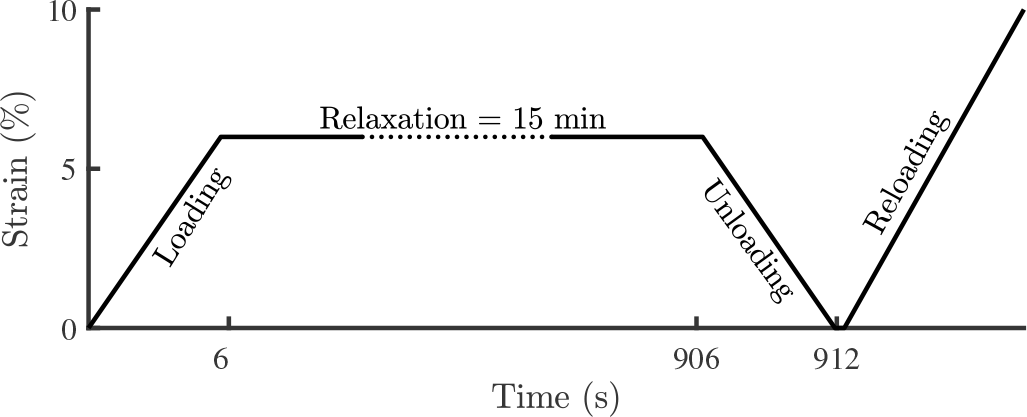
Micro-tensile mechanical testing profile. (A) The samples were ramped to a target grip strain (4%, 6%, or 8%) at a 1%/sec rate (loading phase), held for 15 min (relaxation phase), unloaded at 1%/*sec* (unloading phase), and reloaded to failure (reloading phase). The profile shown corresponds to a 6% target strain test.

### 2.4 Model parameter identification

The models were implemented using a custom written Matlab package for reactive inelasticity (ReactiveBond v1.1 [44]). For parameter identification, we used a constrained multivariable nonlinear optimization method (*fmincon*, Matlab). The root-mean square error (RMSE) was used as the optimization cost function. To minimize the chance of convergence to local minima, we ran the optimization for each of the experimental stress curves using 48 randomly generated initial guesses and the solution with the minimum final residual value was selected as the optimal answer (*MultiStart*, Matlab).

### 2.5 Data analysis

The optimized curve-fit results were individually plotted for each phase of the test, and the resulting model parameters were illustrated using parallel coordinate plots (PCP) to visualize the fit parameters in a compact way [45]. In addition to the individual fit parameters, the means, medians, and the interquartile range (IQR) were calculated and displayed in the PCP. The resulting fit parameters’ similarity to normal distributions were tested using a Jarque-Bera test (*jbtest*. Matlab) with a significance level set at (*p* < 0.05).

The goodness of the fits was assessed using the percent-error between the fits and the experiments, calculated as *%err* = 100(*T*_*fit*_ − *T*_*exp*_)/*T*_*exp,max*_. The IQR of %err is evaluated for both of the models, and plotted for each phase of the deformation as a function of the normalized time of the phase defined as *t* = (*t* − *t*_1_)/(*t*_2_ − *t*_1_), where *t*_1_ and *t*_2_ correspond to the time at the beginning and the end of the phase, respectively. Normalization of time was performed to visualize the patterns of the errors due to variability in strains reached at each phase.

The constitutive relations used for formative bonds were the same between the models, so we hypothesized that the formative bond parameters are not sensitive to the choice of other bond types. To test this hypothesis, we compared the pairs of formative bond model parameters by using Wilcoxon signed rank test (*signrank* Matlab, *p* < 0.05).

### 2.6 Quantification of inelastic effects

The accumulation of inelastic effects are plotted for each bond type in each model. The inelastic effect of damage is represented by the damage parameter *D*_γ_ for formative bonds and permanent bonds. To calculate the inelastic effect of sliding on stress, analogous to the damage parameter, we defined a normalized representation of the sliding effect (when the sliding is occurring) as 
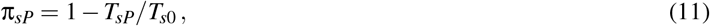
where *T*_*s*0_ is the stress response of sliding bonds using the same intrinsic hyperelasticity relation with no sliding (equivalent to a permanent bond response). As a result, π_*sP*_ takes a value between zero and one, where *π*_*sP*_ = 0 indicates that there is no sliding in the system, and *π*_*sP*_ = 1 has a zero-stress response. To compare the inelastic effect between the models, the median stretches at the 50% inelastic effect (*D*_*fp*_,π_*sP*_,*D*_*fD*_,*D*_*pD*_ = 50%) were compared. Using paired comparison, median stretch of the formative bonds of two models were compared to each other, and the median stretches for sliding bonds were compared to permanent bonds (*signrank*, *p* < 0. 05).

### 2.7 Model validation

To validate the models, we compared the response of both models to the constant strain-rate ramp experimental tests by Szczesny and co-workers [39]. This type of mechanical testing is widely used for material characterization, thus it is a suitable choice for validation [7, 46]. It is important to note that the validation data was not included for the curve-fitting. We used the non-stained constant-rate ramp data from [39] (n=4, Figure 6 in [39]). The median values from the fit results of the models are used for predicting the mechanical response to a 1%/sec deformation curve, up to the peak stress (Table A1). The modulus and the peak stress are compared between the experimental data and the model predictions.

## 3 Results

Both the plastic deformation and damage models resulted in excellent fits to the experimental data (Fig. 3). In particular, both of the models were capable of fitting the loading phase that showed stress stiffening (toe-region) followed by the linear-region (Fig. 3A,E). During the sustained loading of the relaxation phase, stress relaxation was closely fit as a result of using the nth-order kinetics relation for the formative bonds (Fig. 3B,F). The unloading phase had a different nonlinear response compared to the loading phase due to the inelastic behaviors (Fig. 3C,G). The reloading curves showed significant softening before reaching the peak stress that was captured by both models (Fig. 3D,H).

**Fig. 3.**
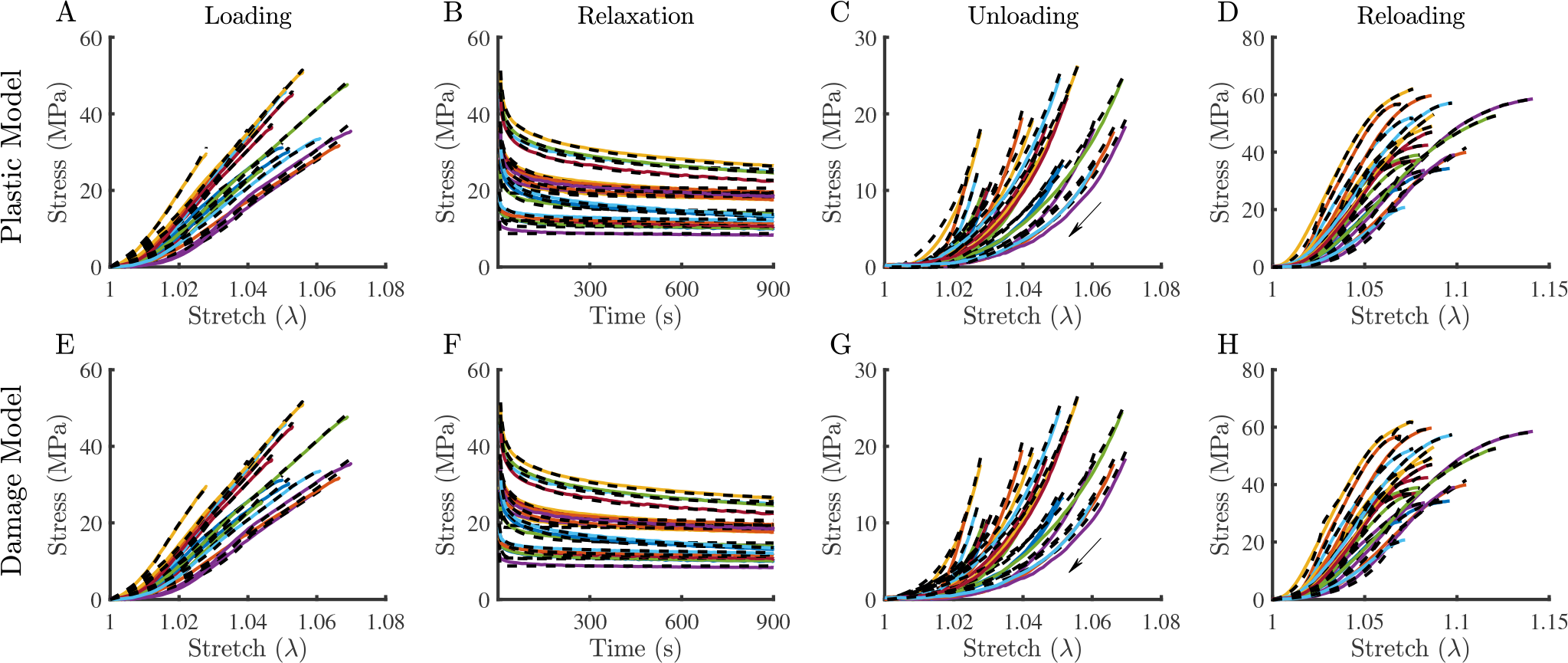
Fit results for individual experiments. The fits are overlaid (black dashed line) on top of individual experimental stress responses, for (A,E) loading, (B,F) relaxation, (C,G) unloading, and (D,H) reloading phases of the experiment. (A-D) are fits to the plastic model and (E-F) to the damage model.

The errors were generally small between the model fits and the experiments, where the median of errors did not exceed 5% (Fig. 4). However, the performance of the models were not equal in all of the phases. In particular, both of the models overestimated the loading toe-region in the loading phase (Fig. 4A,E). The error during relaxation was maximum at the higher stress values-the start of relaxation-and it declined as the relaxation progressed (Fig. 4B,F). Both of the models had minimal overestimation during the unloading phase, where the plastic model had a slightly better performance towards the end of unloading (Fig. 4C,G). Upon reloading, the models underestimated the stress in the nonlinear stiffening region, and the error was lesser at the end of the reloading phase (Fig. 4D,H).

**Fig. 4.**
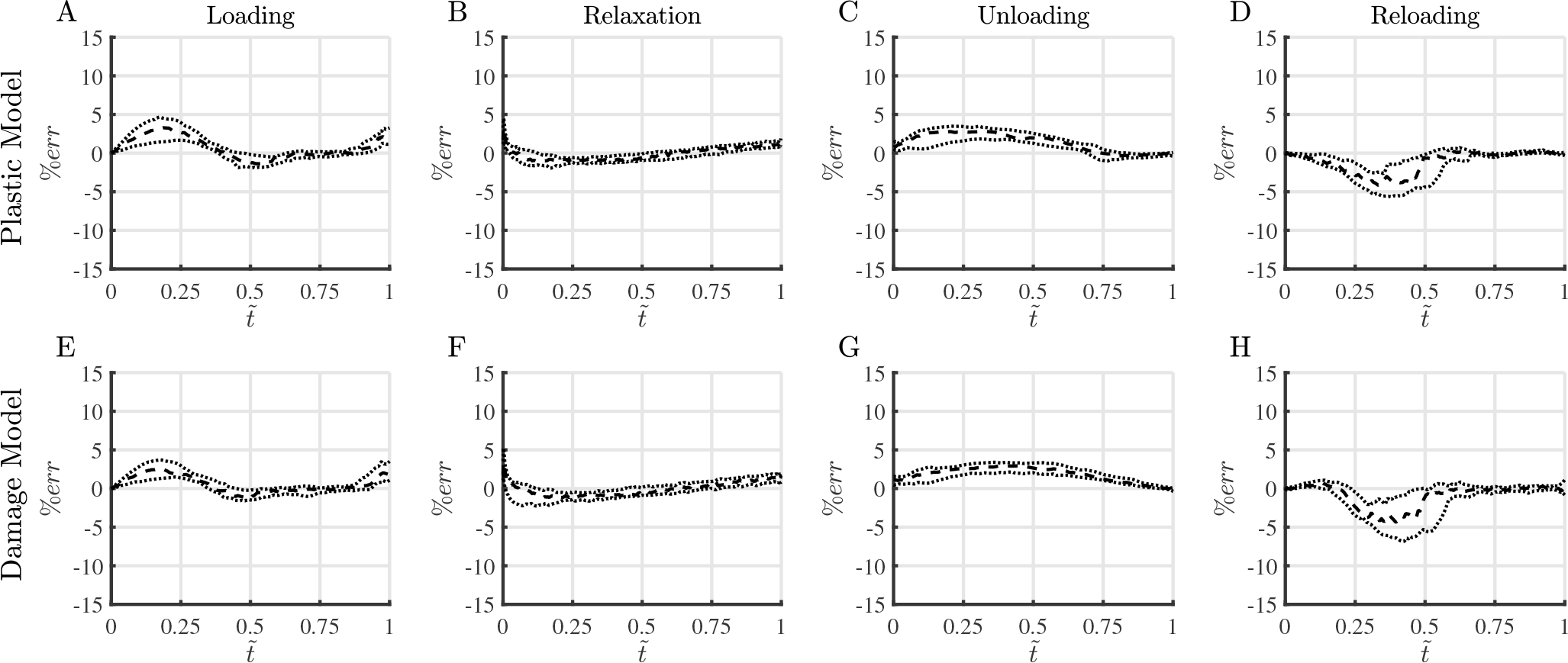
**The IQR of** %*err* **of the model fits** of (A,E) loading, (B,F) relaxation, (C,G) unloading, and (D,H) reloading phases as a function of the normalized time of each phase defined as 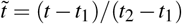, where *t*_1_ is the beginning of the phases and *t*_2_ corresponds to end of the phase. (A-D) are %*err* for the plastic model and (E-F) for the damage model (median = dashed line, *Q*1, *Q*3 = dotted lines).

The fit parameters were visualized using parallel coordinates plots (PCP), where each piece-wise line corresponds to the fit parameters of one sample (Fig. 5). Several model parameters had a different median and mean values, which is a sign for non-normality of the distributions; this was also confirmed by the normality tests that indicates for an overall response the median is more appropriate compared to the mean value (Table A1). Both of the models showed a similar trend in fit parameters for formative bonds, which were modeled using the same parameter sets, but independently fit using each model (Fig. 5). However, the statistical comparisons indicated that formative bonds’ parameters had some differences. In particular, for kinetics of bonds *K*_*f*_ parameters were slightly different (*p* < 0.05), where for plastic model *K*_*f*_ = 0.32*s*^−1^, and for the damage model *K*_*f*_ = 0.34*s*^−1^, but there was no difference between the order of the kinetics relation (*N*_*f*_ ~ 1.5). Additionally, contrary to our hypothesis all of the pairs of intrinsic hyperelasticity (*C*_1*f*_, *C*_2*f*_) and damage (*k*_*f*_, *I*_*f*_, (*r*_0_)_*f*_) parameters were different, except for only *I*_*f*_ that did not show a difference (*p* < 0.05) (Table A1). The sliding parameters of sliding bonds and the damage parameters of the permanent bonds were dissimilar, which was expected since they belong to different bond types (Figure 5 and Table A1).

**Fig. 5.**
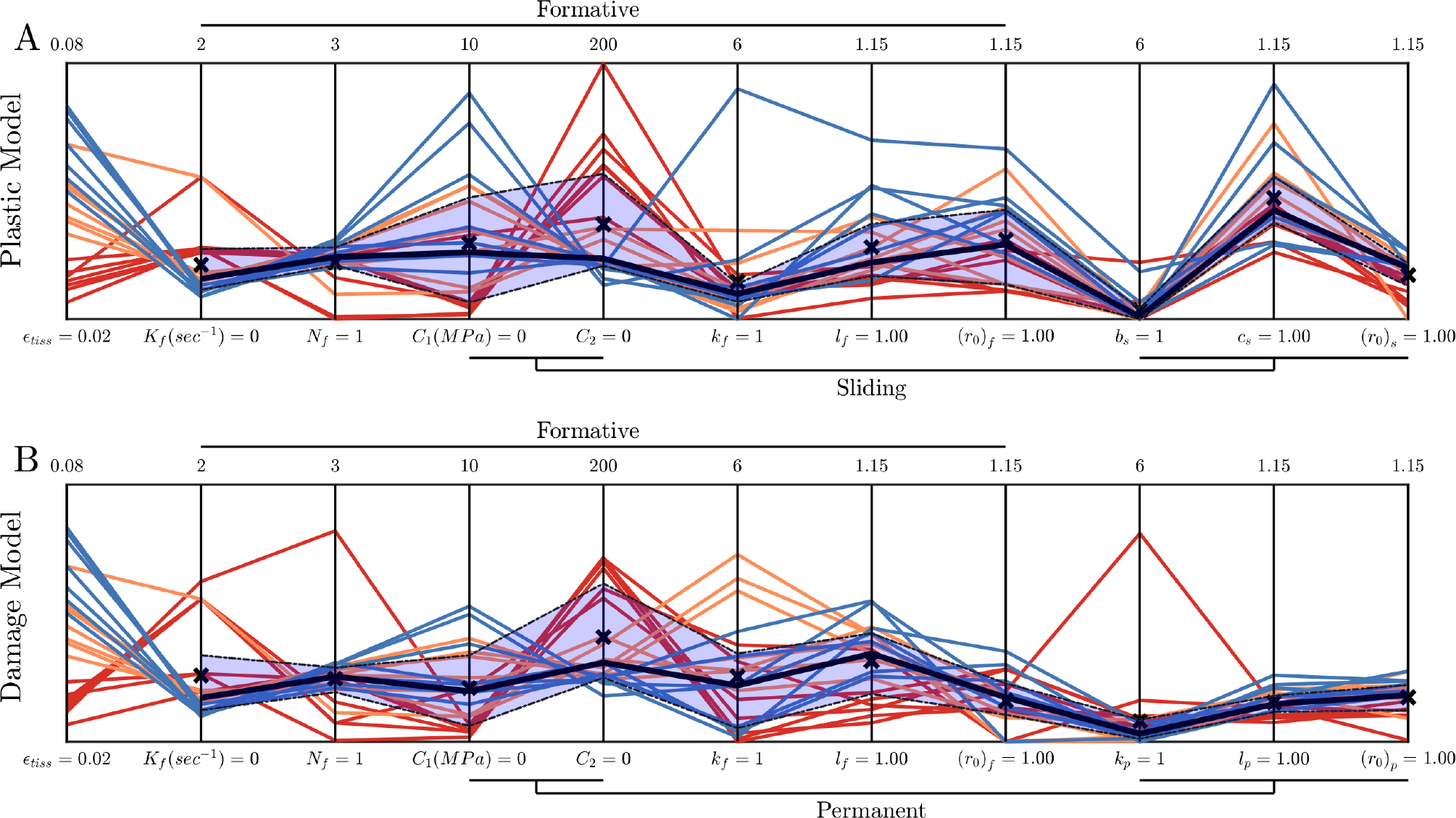
**Parallel coordinates plots of the fit parameters** for (A) plastic deformation model and (B) damage models, color-coded based on the nominal grip strain (red = 4%, orange = 6%, and blue = 8%). The interquartile range (IQR) is shaded, where the median value is plotted using a solid black line. The mean value is marked with “×”.

The inelastic effects on the bonds were calculated for each experiment and the stretch at which 50% inelastic effect for a bond type was reached (Fig. 6). In the plastic model, the formative bonds’ damage reached *D*_*fP*_ = 0.50 at the stretch value of 1.07[1.04,1.09] (read as *median*[*Q*1, *Q*3], Fig. 6A). The sliding bonds in the plastic model, π_*sP*_ reached 50% at the stretch of 1.05[1.04,1.06] (Fig. 6B). For the damage model, the formative bonds reached *D*_*fD*_ = 0.50 at λ = 1.06[1.05,1.07], which had a slightly different distribution compared to that of formative bonds in the plastic model (*p* < 0.05) (Fig. 6C) and the permanent bonds *D*_*pD*_ = 0.5 was reached at λ = 1.05 [1.03,1.05] which was similar to the sliding bonds (Fig. 6D). When looking at the individual samples, due to the non-recoverable nature of the inelastic effects during reloading, higher values were reached for all of the inelastic parameters compared to their counterpart in the loading phase (Fig. 6). In addition to the inelastic effects, for sliding bonds we plotted the reference stretch λ_*s*_ that reached the small value of 1.01 [1.0,1.02] during loading, and it increased to 1.05[1.04,1.05] at the end of reloading phase (Fig. 6B).

**Fig. 6.**
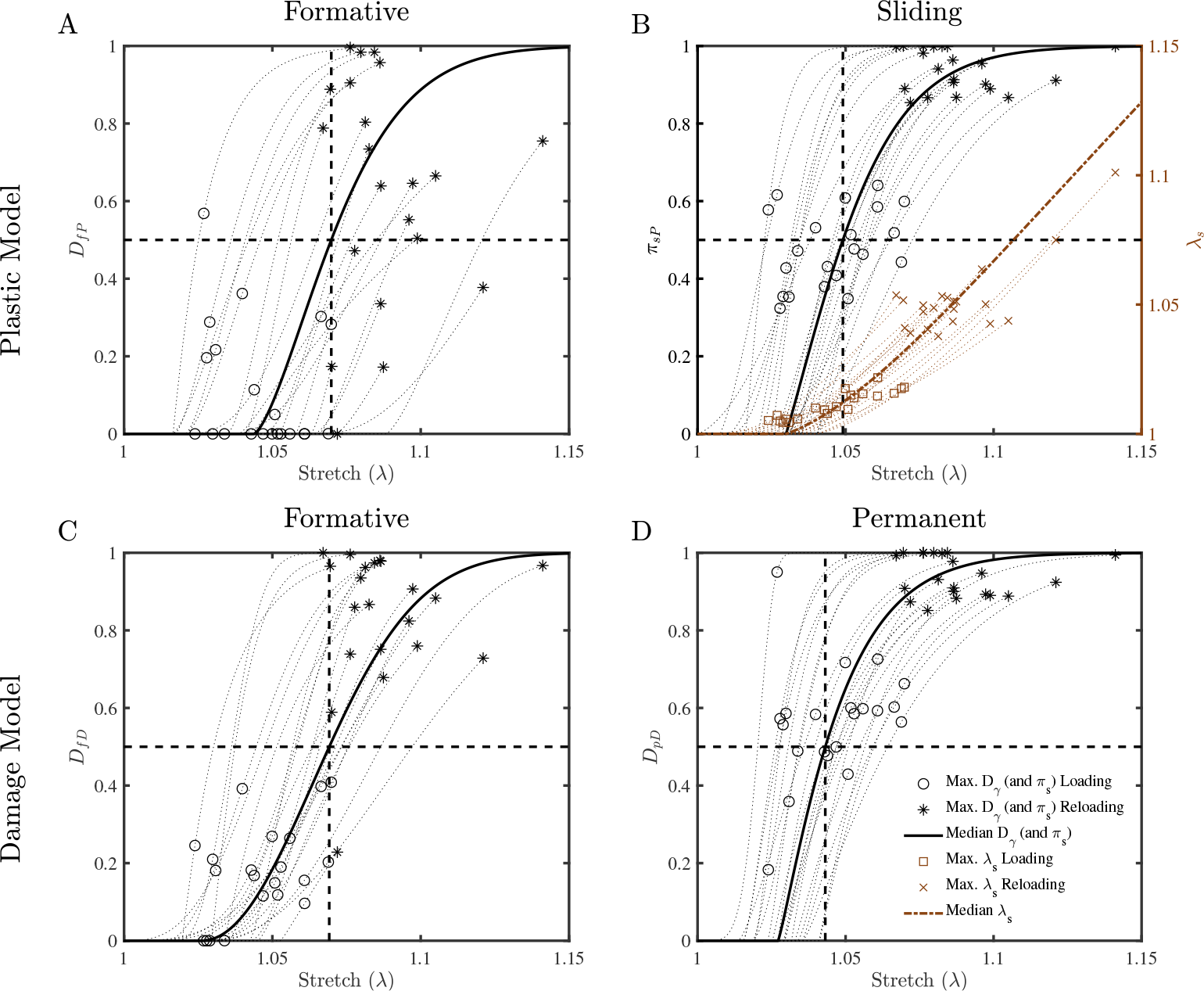
Accumulation of damage and plastic deformation. The overall response for accumulation of damage (*D*_γ_), and normalized palstic deformation (*π*_*s*_) are plotted for each bond type using the median of the fit parameters (solid line). The 50% inelastic effect lines are marked using horizontal straight lines and the median stretch of all samples’ intersected with this 50% inelastic efect line is marked with a vertical line. Additionally, the maximum damage and normalized plastic deformation reached during individual fits in loading (open circle), and reloading (star) phases for the individual fits. For the sliding bonds, the sliding stretch (λ_*s*_) is also plotted (brown dashed line) and the maximum plastic deformation stretch during loading (open square) and reloading (cross) phases is plotted with respect to the right y-axis (brown).

To validate the model fits, the median parameter from both of models were used to predict independent experiments, where both models produced similar stress-strain curves that include the initial stiffening (toe-region) and subsequent increase in stress followed with softening (Fig. 7). Average experimental modulus and peak stress were 909 ± 27 MPa and 58 ± 4 MPa, respectively [39]. The predicted modulus of the plastic model was 832 MPa and the peak stress was 31 MPa. Respectively, these values were −8% and −46% different from the experiments. For the damage model, the modulus was 1036 MPa, and the peak stress was 64 MPa, which were 14% and 10% different from their experimental counterparts, respectively. In overall, both models produced similar stress-strain curve shapes, and modulus values; however, the damage model resulted in a closer peak stress prediction when compared to the plastic model (Fig. 7).

**Fig. 7.**
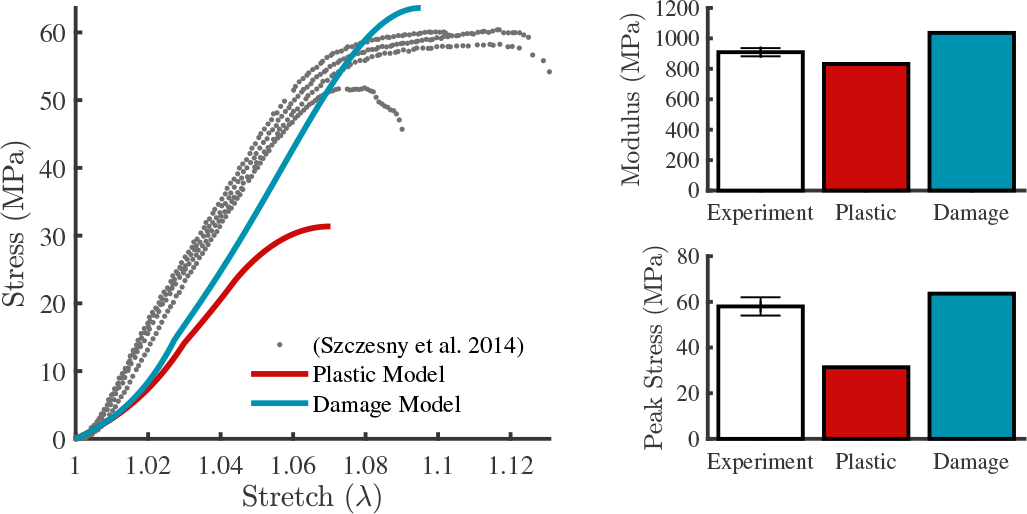
Validation plots for prediction of plastic and damage models showing the comparison of the experimental data from [39] to the predictions of the plastic model (red) and damage model (blue) to a ramp loading.

## 4 Discussion

### 4.1 Comparison between the plastic deformation and damage models

In this study we successfully applied the theoretical framework of reactive inelasticity to tendon experimental data. We implemented two independent models, specific to plastic deformation or damage, and showed excellent fits to the experiments (Fig. 3), and visualized the outcome parameters using parallel coordinate plots, which enhances visualization of the complex relationships among them. The fits showed that the plastic model had slightly smaller errors towards the end of the unloading phase, which was due to inclusion of plastic deformation that agrees with previous studies [13,14]. However, when comparing the model predictions using the resultant fit parameters to independent validation experimental data, both of the models predict similar stress-strain curves, with the damage model having more similar behavior (Fig. 7). These results do not strongly favor one approach over the other, and both of the inelastic mechanisms may contribute to the mechanical response, thus further investigations and different experimental testing protocols are needed to differentiate between tendon plastic deformation and damage behaviors.

### 4.2 Interpretation of inelastic effects

The J-shaped form for the accumulation of inelastic behaviors in our study (Fig. 6) are consistent with the findings of a recent study that reported the increase in denatured collagen during tensile loading by using collagen hybridizing peptides (CHP) [20]. This indicates that the accumulation of inelastic effects may be correlated with collagen denaturation due to mechanical loading. However, the CHP study suggested onset of damage at higher strains (~ 8% strain) [20] compared to our study ((*r*_0_)_γ_ < 5% Table A1) and other studies [13,14,18]. This suggests that in addition to collagen denaturation, other molecular mechanisms and proteins in extra-cellular matrix (perhaps elastin or decorin) may play a role in the inelastic mechanical response of tendon [47–49]. While identification of the relationships between tendon’s inelastic mechanics and molecular disruptions require further investigations, the inelastic effects can be quantified by using mechanical modeling for understanding the mechanical properties of tendon, and assessing its altered mechanical properties in various stages of disease, injury, and healing [50–52].

### 4.3 Remarks about RIE modeling

We hypothesized that the formative bond parameters would be the same for both models regardless of the choice of sliding or permanent bonds for the equilibrium response. While they were similar, there were some unexpected and statistically significant differences (Table 1). This may be due to the simplifying assumption we made that the formative bonds had the same intrinsic hyperelasticity parameters as the sliding or permanent bonds (Table 1). We made this assumption in order to use the minimum number of variables to model the tendon response. It is unclear if full independence of bond parameters would provide a benefit over the current modeling framework. Nonetheless, it is likely that including interaction terms between bonds or addition of more bonds and parameters, as a spectrum [53,54], could be beneficial for accurate modeling of tissue’s inelastic behaviors.

### 4.4 Sources of error

The percent-error (%err) of the models were quite small, but both of the models had relatively large errors in the toe-regions and the beginning of the relaxation phase (Fig. 4). Smoother fits could be achieved with more elaborate constitutive relations, such as fiber-recruitment for intrinsic hyperelasticity [55, 56] or normalized energy and stress for sliding and damage [34, 57, 58], and by adding derivatives of the stress response to the cost-function. For the relaxation phase, despite some deviations at the beginning, this phase was well fit by using a generalized nth-order kinetics for formative bonds (Fig. 4B,F). The error in the relaxation phase may be related to fluid flow dependent viscoelasticity [43,59], which was not included in this study. We expect this to be a small effect for tail tendon; however, it can be added by using a biphasic mixture with RIE as the solid phase [30,60]. Additionally, we assumed that the kinetics rate of bond breakage and reformation is not dependent on the level of strain; however, adding strain-dependence to kinetics parameters might be necessary for modeling more complicated loading scenarios such as incremental stress-relaxation [24,61].

### 4.5 Conclusion and future direction

In conclusion, we applied the theoretical framework of reactive inelasticity (RIE) to model viscoelasticity, plastic deformation, and damage to determine tendon’s inelastic mechanical response. This study is novel in that (1) it investigated tendon inelastic effects without a prior assumption about the softening mechanism and compared the two modeling approaches for inelasticity, (2) it demonstrated numerical values for inelasticity effects on tendon mechanical response, and (3) it provides a path forward to relate the molecular structure of tendon to its mechanical response. We applied the models to the experimental data, and validated the model by comparing the predictions to an independent set of experimental data. The both models were successful in fitting and predicting the experimental data, although the plastic model had slightly better fits during the unloading phase, and the damage model had better predictions of the independent validation data set. However, the results largely indicate that deformation-driven experiments can be equally described with both plastic deformation and damage perspectives. Thus, distinguishing plastic deformation and damage mechanisms will require further experimental studies using different loading protocols.

**Table A1.** Fit parameter results

|  | Bond Type | Kinetics | Intrinsic Hyperelasticity | Sliding | Damage |
| --- | --- | --- | --- | --- | --- |
| Plastic Model | **Formative** | $[K(s^{-1}), N]_{fP}$ | $[C_1(\text{MPa}), C_2]_{fP}$ | - | $[k, l, r_0]_{fP}$ |
|  | Mean | [0.43*, 1.44*] | [2.98, 74.34*] | N/A | [1.74*, 1.04*, 1.05] |
|  | Median | [0.32<sup>a</sup>, 1.49] | [2.62<sup>b</sup>, 47.47<sup>c</sup>] | N/A | [1.50<sup>d</sup>, 1.03, 1.04<sup>e</sup>] |
|  | Q1 | [0.23, 1.42] | [0.67, 41.76] | N/A | [1.34, 1.03, 1.02] |
|  | Q3 | [0.55, 1.56] | [4.76, 113.87] | N/A | [1.70, 1.06, 1.06] |
| | **Sliding** | - | $[C_1(\text{MPa}), C_2]_{sP}$ | $[b, c, r_0]_{sP}$ | - |
|  | Mean | N/A | *same as fP* | [1.18*, 1.07*, 1.03] | N/A |
|  | Median | N/A |  | [1.04, 1.06, 1.03] | N/A |
|  | Q1 | N/A |  | [1.00, 1.06, 1.02] | N/A |
|  | Q3 | N/A |  | [1.22, 1.08, 1.03] | N/A |
| Damage Model | **Formative** | $[K(s^{-1}), N]_{fD}$ | $[C_1(\text{MPa}), C_2]_{fD}$ | - | $[k, l, r_0]_{fD}$ |
|  | Mean | [0.52*, 1.49*] | [2.09, 81.48] | N/A | [2.27, 1.05, 1.02] |
|  | Median | [0.34<sup>a</sup>, 1.51] | [1.97<sup>b</sup>, 61.53<sup>c</sup>] | N/A | [2.10<sup>d</sup>, 1.05, 1.03<sup>e</sup>] |
|  | Q1 | [0.25, 1.38] | [0.61, 50.01] | N/A | [1.26, 1.03, 1.02] |
|  | Q3 | [0.67, 1.58] | [3.37, 123.12] | N/A | [2.72, 1.06, 1.03] |
| | **Permanent** | - | $[C_1(\text{MPa}), C_2]_{pD}$ | - | $[k, l, r_0]_{pD}$ |
|  | Mean | N/A | *same as fD* | N/A | [1.40*, 1.02, 1.03] |
|  | Median | N/A |  | N/A | [1.16, 1.02, 1.03] |
|  | Q1 | N/A |  | N/A | [1.06, 1.02, 1.02] |
|  | Q3 | N/A |  | N/A | [1.43, 1.03, 1.03] |

\* The distribution is non-normal ( $p < 0.05$ )
a-e The medians of the parameter distributions are different ( $p < 0.05$ )

## Acknowledgment

We would like to thank Prof. Ryan Zurakowski, University of Delaware, for donation of computational resources. Research reported in this publication was supported by the National Institute of Biomedical Imaging and Bioengineering of the National Institutes of Health under award number R01EB002425. The content is solely the responsibility of the authors and does not necessarily represent the official views of the National Institutes of Health.

## Appendix: Fit parameters

The detailed descriptive statistics of the fit parameters (Table A1). The normality tests are marked on the mean values and the comparison between the formative bond parameters are marked on the median values.

